# Identification of putative GATA3 regulatory elements and comparison of GATA3 distribution in cochleae of mice, rats, macaques, and humans

**DOI:** 10.1101/2022.10.12.511664

**Authors:** Sumana Ghosh, Robert Wineski, Ivan A. Lopez, Akira Ishiyama, Punam Thapa, Bradley J. Walters

**Affiliations:** Department of Otolaryngology – Head and Neck Surgery, University of Mississippi Medical Center, Jackson, MS 39216, U.S.A.; Dept. of Head & Neck Surgery, David Geffen School of Medicine, University of California at Los Angeles, Los Angeles, CA 90095, U.S.A.

**Keywords:** auditory, vestibular, cochlea, spiral ganglion, regeneration, development, hair cells

## Abstract

The transcription factor GATA3 plays a critical role in the development of neurons and sensory epithelia of the inner ear. In mouse cochleae, GATA3 is downregulated in certain supporting cells (SCs) and in type I spiral ganglion neurons (SGNs) after development. This reduction of GATA3 in SCs severely limits *Atoh1*-induced hair cell (HC) regeneration and suggests that a similar downregulation in human cochleae may be limiting for regenerative therapies. However, it is unknown whether GATA3 is similarly or differentially regulated in primates versus rodents. Using CAGE-seq data, we compared over 40 putative *GATA3* regulatory elements across species and found both conserved and non-conserved sequences. To assess whether cochlear GATA3 distribution is similar or different between rodents and primates, we immunostained cochleae from mice, rats, macaques, and humans using antibodies raised against highly conserved GATA3 peptide sequences. GATA3 immunostaining in the organs of Corti from all four species revealed a large degree of conservation, where SCs medial and lateral to cochlear HCs exhibited robust nuclear GATA3 immunolabeling, but pillar and Deiters cells had significantly reduced GATA3 immunoreactivity. In all four species, GATA3 was expressed in a subset of SGNs that largely co-expressed peripherin suggesting they were type II SGNs. Only one difference emerged, wherein human cochlear inner hair cells were not GATA3 immunoreactive despite being so in the other species. Overall, the pattern of GATA3 expression in primates appears similar to rodents and reinforces the notion that ATOH1 mediated regenerative therapies may be limited by reduced GATA3 expression in adult SCs.

## INTRODUCTION

GATA binding protein 3 (GATA3) is a zinc finger transcription factor with a wide distribution of expression in mammalian species and is important in the development and function of many different tissues and cell types. The importance of GATA3 is highlighted by the deficits observed in HDR syndrome which is primarily caused by a haploinsuficency of GATA3 and results in hypoparathyroidism, renal disease, and sensorineural hearing loss (Van Esch & Devriendt, 2001) During mouse inner ear development, GATA3 is expressed in the otic vesicle and the immature cochleovestibular ganglion (Lawoko-Kerali et al., 2002; Lillevüli et al., 2004; Rivolta & Holley, 1998). Specifically, GATA3 expression defines the otic epithelium during invagination of the otic cup at embryonic day (E) 8.5, continues to be expressed throughout much of the otic vesicle and gradually becomes restricted to the delaminating neuroblasts (∼E11.5) and the prosensory domains that will give rise to the hair cells (HCs) and supporting cells (SCs) of the auditory and vestibular epithelia. GATA3 is subsequently downregulated from the vestibular tissues, but continues to be expressed in the organ of Corti and SGN of the cochlea. In the organ of Corti, GATA3 appears to be further downregulated in outer HCs (OHCs) by postnatal day 0 (P0), but is still expressed in all of the SCs and in the inner HCs (IHCs) (Walters et al., 2017). Genetic ablation of *Gata3* during embryogenesis reveals that it is critical for the development of the sensory structures of the inner ear with particular importance to the cochlear epithelium and innervation thereof (Duncan et al., 2011; Haugas et al., 2012). *Gata3* has also been shown to be a critical co-factor for *Atoh1* (Masuda et al., 2012) and possibly *Eya1* (Duncan & Fritzsch, 2013), and is therefore important for the process of HC differentiation. Insufficiency of *Gata3*, as in animal models of HDR syndrome, show that it dramatically affects SGN survival (Luo et al., 2013), HC survival (Van Der Wees et al., 2004) and HC function (Van Looij et al., 2005).

In studies using mouse models, it has been reported that the pattern of GATA3 expression continues to be refined postnatally as the cochlea matures and the animals acquire the ability to hear. Beginning late in embryonic development and proceeding into adulthood, GATA3 declines to levels undetectable by immunostaining in type I SGNs, cochlear OHCs, pillar cells (PCs), and Deiters’ cells (DCs) (Nishimura et al., 2017; Walters et al., 2017). While neither the functional effects of these postnatal changes, nor the roles of persistent GATA3 expression in cochlear IHCs and non-pillar or non-Deiters’ SCs, are well-elucidated, the downregulation of GATA3 in the PCs and DCs has been shown to be a limiting factor for the forced conversion of these SCs to HCs via ectopic expression of *Atoh1* in the cochleae of young adult mice (Walters et al., 2017).

This necessity of *Gata3* for *Atoh1*-induced cellular conversion is important for therapeutic objectives in that the two most prominent methods currently being pursued to elicit HC regeneration are 1) the direct ectopic expression or manipulation of *Atoh1*, or 2) the inhibition of Notch signaling, which acts via the upregulation of *Atoh1* (Izumikawa et al., 2008; Mizutari et al., 2013; Walters et al., 2017; Yamamoto et al., 2006). Recent work has shown that GATA3 is a co-factor that forms a transcriptional activation complex with ATOH1 to upregulate HC-specific genes (Masuda et al., 2012). Furthermore, loss of GATA3 from PCs and DCs correlates with an inability to regenerate mammalian cochlear hair cells via ectopic *Atoh1* expression, and this failure of *Atoh1* to convert PCs and DCs to HC-like cells can be overcome by ectopic expression of *Gata3* in PCs and DCs (Walters et al., 2017). Thus, it would seem that the loss of GATA3 with age in a large subset of SCs might prove a hinderance to either Notch or Atoh1 based approaches to regenerate HCs for therapeutic benefit. As both strategies are currently being actively pursued in clinical and preclinical trials, we sought to determine whether the pattern of GATA3 downregulation in certain SCs with age was specific to mice, or if it was conserved between rodents and human and non-human primates.

Indeed, in recent years, many findings from rodents have failed to be translated to humans, in part, due to differences in gene regulation and expression (Hackam & Redelmeier, 2006). Also, in recent years, several papers have highlighted differences in the patterns of expression of several proteins known to be important in inner ear function when comparisons are made between primate and rodent species (Hosoya, Fujioka, Ogawa, et al., 2016; Hosoya, Fujioka, Okano, et al., 2016; Matsuzaki et al., 2018). Furthermore, in the case of *GATA3*, several regulatory DNA elements have been identified, some of which appear to be conserved in mice and humans, and others that are not (Hasegawa et al., 2007; Ohmura et al., 2016).

While a cis regulatory sequence dubbed the “Otic-specific enhancer” of *Gata3* has been identified (Mariguchi et al 2018), it is not clear whether this element regulates Gata3 expression only during development or if perhaps other regulatory sequences are responsible for the pattern of Gata3 expression seen in the adult inner ear. Therefore, we sought to further investigate putative regulatory elements of GATA3 as well as the distribution of GATA3 expression in the cochleae of adult rats, mice, macaques, and humans. Our analysis suggests that there are many potential GATA3 regulatory elements and that several of these are not conserved between rodents and primates. However, the pattern of GATA3 expression in the adult organ of Corti does appear to be largely conserved, with only one obvious difference, a lack of GATA3 immunoreactivity in human inner HCs. Importantly, GATA3 is not highly expressed in PCs and DCs of any of the four species in this study, including humans, suggesting that these cells may respond similarly to mouse PCs and DCs when ectopic *ATOH1* is applied and may therefore largely fail to convert into HC-like cells.

## MATERIALS and METHODS

### Identification and comparison of potential GATA3 regulatory elements

Putative *GATA3* promoter and enhancer sequences were identified using the human promoters selector and the human enhancer selector tools on SlideBase (http://slidebase.binf.ku.dk/) and the FANTOM5 consortium data (Andersson et al., 2014; Forrest et al., 2014; Ienasescu et al., 2016). Searches for GATA3 resulted in the identification of thirty-two non-contiguous potential promoter sequences and ten potential enhancers. These were then queried against the published genomic sequences for *Mus muscularis* (NC_000068.7, GRCm38.p6)(Church et al., 2009, 2011; Waterston et al., 2002), *Rattus norvegicus* (NC_005116.4, Rno6.0)(Gibbs et al., 2004), and *Macaca fascicularis* [NC_022280.1, Macaca_fascicularis_5.0 (Higashino et al., 2012)] using the NCBI Blastn tool (Altschul et al., 1990; McGinnis & Madden, 2004; NCBI, 2015). If no matching sequence could be identified on the appropriate chromosome by Blastn, the genomic sequences were checked manually using text find searches and the homologous regions were then directly compared using Clustal-omega (Daugelaite et al., 2013; Sievers et al., 2011). A thorough literature search was also conducted to obtain regulatory sequences which have been previously identified and validated experimentally (Supplemental table S1). These sequences were then compared between mouse and human genomes using the approach described above. Peptide sequences for GATA3 in human, macaque, rat, and mouse were obtained from the NCBI nucleotide database and aligned using Clustal-omega.

### Non-human tissue preparation and immunostaining

All of the mice, rats, and macaques used for this study were housed and cared for according to IACUC approved protocols and NIH guidelines for animal care and use in the Center for Comparative Research at the University of Mississippi Medical Center (UMMC). CD-1 Mice (Charles River laboratories, Wilmington, MA) at either postnatal day (P)0 or P150 and Long Evans rats at P150 (Harlan, Indianapolis, IN) were euthanized by CO2 inhalation and then their temporal bones were removed and immersed in 4% paraformaldehyde (PFA) for 24 hours.

Mouse temporal bones were then decalcified in 0.125 M EDTA for 24-48 hours, while rat temporal bones were decalcified in 0.25 M EDTA for 7 days. The samples were then placed in 30% sucrose in 0.1M PBS and stored at 4°C for 3-4 days before being cryosectioned.

Macaques (between 4-6 years of age) were sedated with ketamine HCl, and then deeply anesthetized with sodium pentobarbital (50 mg/kg, IP). They were then transcardially perfused with 0.1 M, pH 7.2 phosphate buffered saline (PBS), followed by 3 L of a mixture of 1% PFA and 1.5% glutaraldehyde in 0.1 M PBS (1 X PBS, pH 7.2). Temporal bones were then dissected out from the skulls and decalcified in 0.5M EDTA on an orbital shaker at room temperature for 3-4 weeks with EDTA being changed out every 3-4 days. Following decalcification, temporal bones were immersed in 30% sucrose in PBS and stored at 4°C for 7 days before being cut into 14-16 micron thick sections on a cryostat (Leica CM3050). For each species, and for each GATA3 antibody used, sections from the cochleae of at least 4 individuals were immunostained and imaged, and equal numbers of males and females were used. Sections were treated with pH 6.0 antigen unmasking solution (Vector laboratories) for 45 minutes at 98ºC, washed, and then treated with imageIT-Fx signal enhancer for 30min at room temperature. The samples were then washed in PBS, blocked in 1%BSA, 10% normal serum, and 1% Triton X in PBS for 1 hour then incubated in primary antibodies overnight at 4ºC, washed the next day, and placed in secondary antibodies for 1.5 – 3 hours at room temperature. The details of the antibodies used for this study is listed in Table1.

### Tissue preparation and immunostaining of human samples

Celloidin embedded sections of human temporal bones were processed as described previously for immunohistochemistry and brightfield microscopy (Lopez et al., 2016). Briefly, sections were placed in a glass Petri dish and immersed in ethanol saturated with sodium hydroxide solution (1 g/ml of NaOH in ethanol), diluted 1:3 with 100% ethanol for 1 hour, followed by 100% ethanol, 50% ethanol, and distilled water for 10 minutes each. Slides were placed horizontally in a glass Petri dish containing antigen retrieval solution (Vector Antigen Unmasking Solution, Vector Labs, Burlingame, CA diluted 1:250 with distilled water) and heated in the microwave oven using intermittent heating of two 2-minute cycles with an interval of 1 minute between the heating cycles. Slides were allowed to cool for 15 minutes, followed by 10 minutes wash with phosphate buffered saline solution (PBS, 0.1 M, pH 7.4). Sections were blocked in PBS containing 1% bovine serum albumin fraction V (Sigma-Aldrich, St. Louis, MO) and 0.1% Triton X-100 (Sigma-Aldrich) for 1 hour, and incubated subsequently with the antibody against GATA3 in blocking solution (1:100) for 72 hours at 4°C in a humid chamber and processed for IHC as described. Slides were mounted in aqueous mounting media (Fisher Labs). Absorption controls were conducted by preincubating the BD Biosciences primary antibody with equimolar recombinant human GATA3 peptide (MyBioSource #MBS9221421) for several hours prior to the application of the antibody to sections. Controls where only the secondary antibody was used were also performed. The Institutional Review Board (IRB) of UCLA approved this study of human temporal bones (IRB protocol #10-001449, AI) and methods used in this study are in accordance with NIH and IRB guidelines and regulations. Informed consent was obtained from each patient before inclusion in the study of temporal bone sections.

Positive control samples of human tonsil and temporal bone sections from an 8 year old male were obtained from the NIDCD National Temporal Bone Laboratory at UCLA and processed for immunohistochemistry as described previously (Atayar et al., 2005; Lopez et al., 2016)).

For immunofluorescent visualization of GATA3 in human temporal bone sections, adult human temporal bones were resected from cadavers generously donated to the UMMC body donation program. All cadavers were arterially embalmed within 6 hours of death with fluid containing 1.5% formalin, 10% phenol, and 15% glycerin. Cadavers were then maintained in anatomy tables containing 0.1% formaldehyde and 0.5% phenol for 2-3 months and periodically removed from fixative for anatomy education. For the analysis of GATA3 expression in adult tissues, temporal bones from at least 4 different individuals were used (2 male and 2 female) for each GATA3 antibody. All of the individuals were over the age of 50. Temporal bones were resected with an autopsy saw (Stryker Corp., Kalamazoo, MI), and then immersed in 0.5 M EDTA on an orbital shaker for 3 - 4 weeks. Every 3-4 days, decalcified bone was shaved away using a scalpel, and EDTA was replaced. Samples were immersed in 30% sucrose in PBS for 7 days before being cut into 16-20 μm sections on a cryostat. All samples were immunostained as described above for rodents and macaques except the hair cells were labelled with mouse anti myosin VIIa and all secondary antibodies were used at 1:500 dilution. For GATA3 labelling in SGNs, higher percentages of triton-X were used for blocking (2%) and dilution buffer (1%), and for co-labeling with peripherin, samples were pre-treated in TE buffer (pH 9) for 44 minutes at 95°C instead of the low pH antigen retrieval step noted above. All immunolabeled sections were imaged using either a Nikon C2+ confocal microscope or Zeiss LSM880 confocal microscope.

### Antibody Characterization

Characterizations of antibodies that have been accomplished by the manufacturers of commercial antibodies or in previous publications are described in Table 1. In this report, the GATA3 antibody from BD-Biosciences (Mouse, Monoclonal IgG1, catalog # 558686, RRID AB_2108590) was thoroughly tested for specificity (Figure 1). This includes demonstrating positive control staining of inner ear tissues from mice and humans, as well as in human tonsil tissue. For human tissues, absorption controls and secondary antibody-only controls were performed, and in mice, a *Gata3* conditional knockout mouse was used, as negative controls.

**Table 1:**
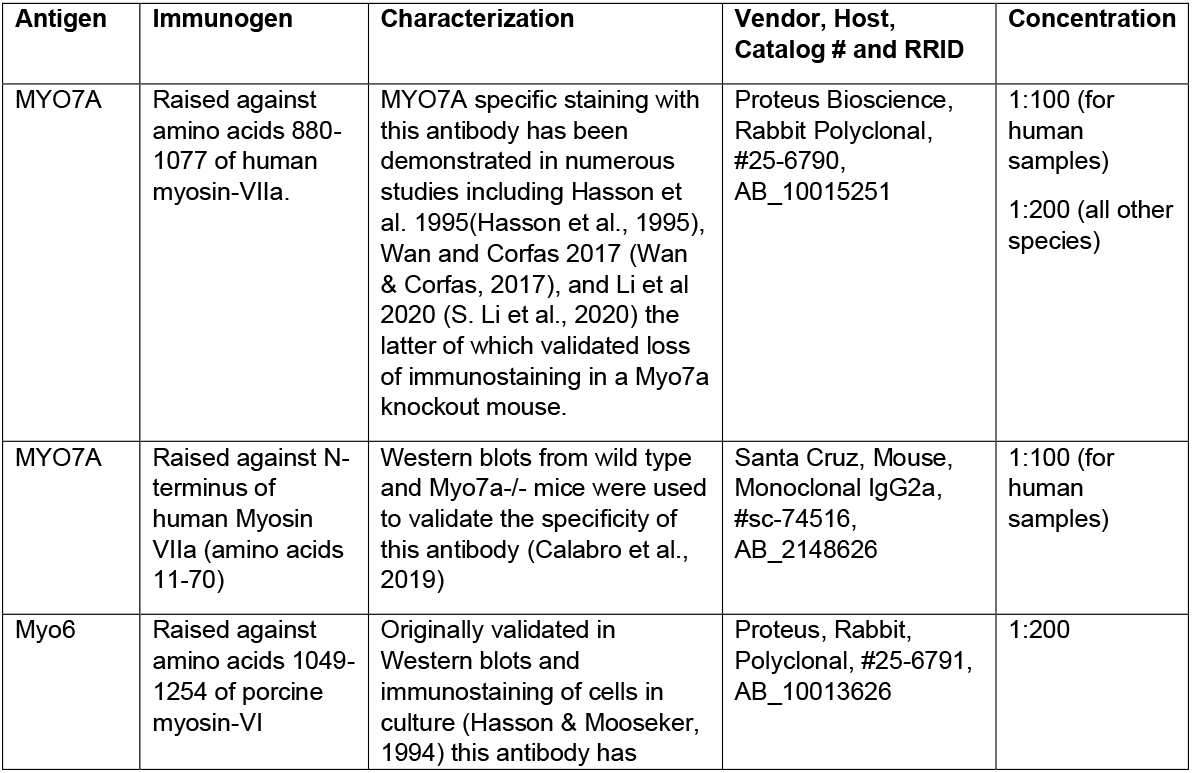
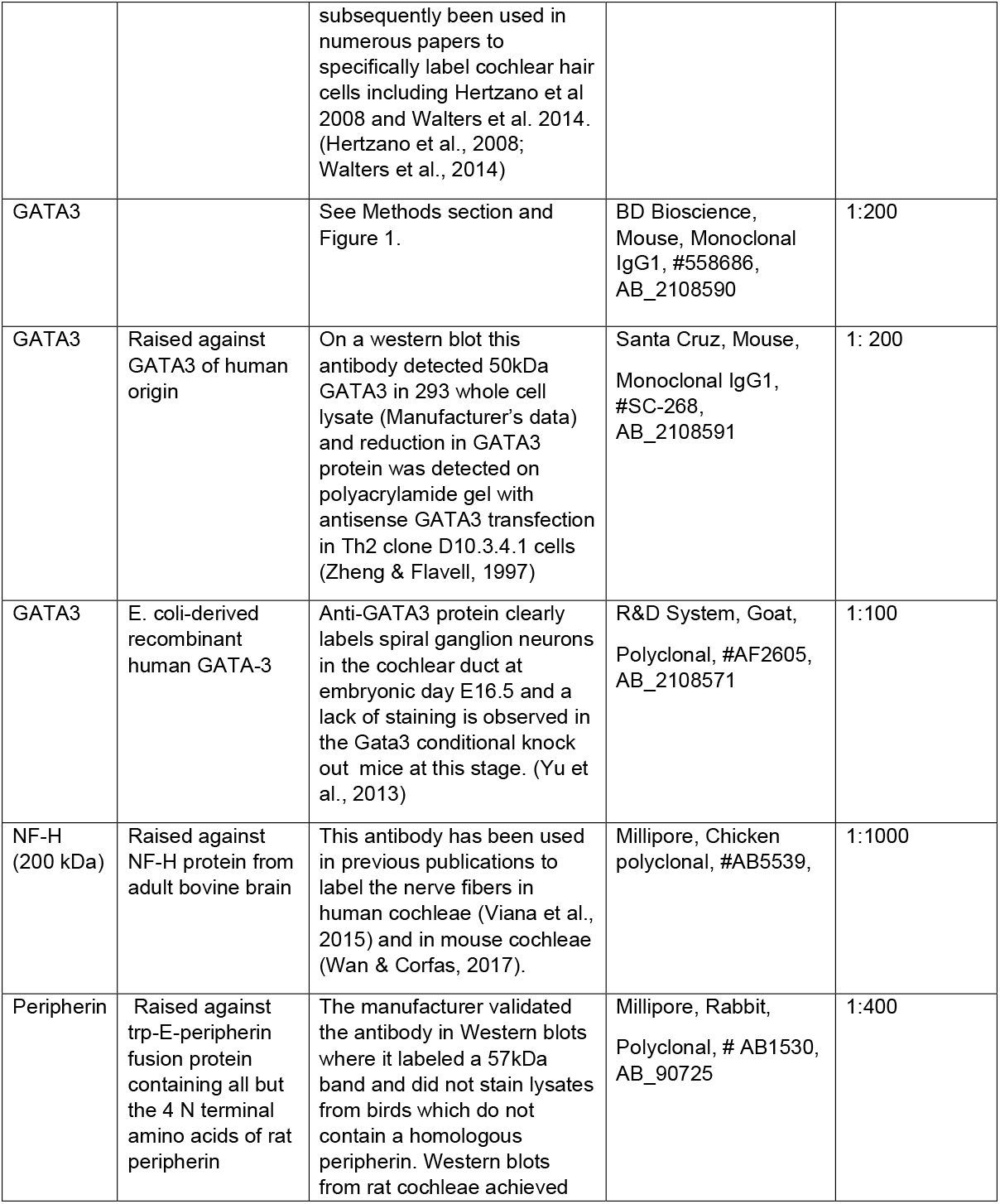
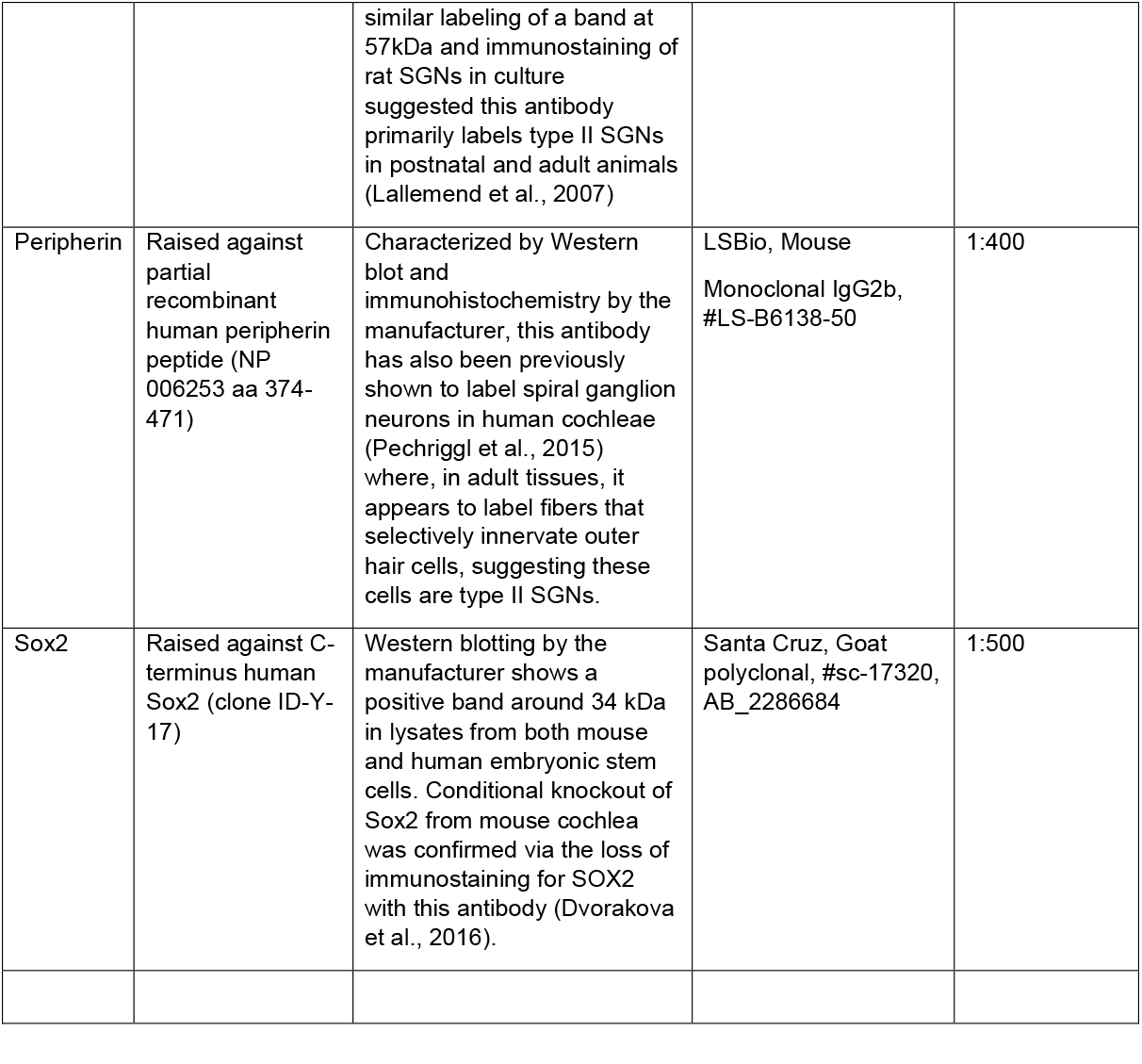
List of Antibodies.

**Figure 1:**
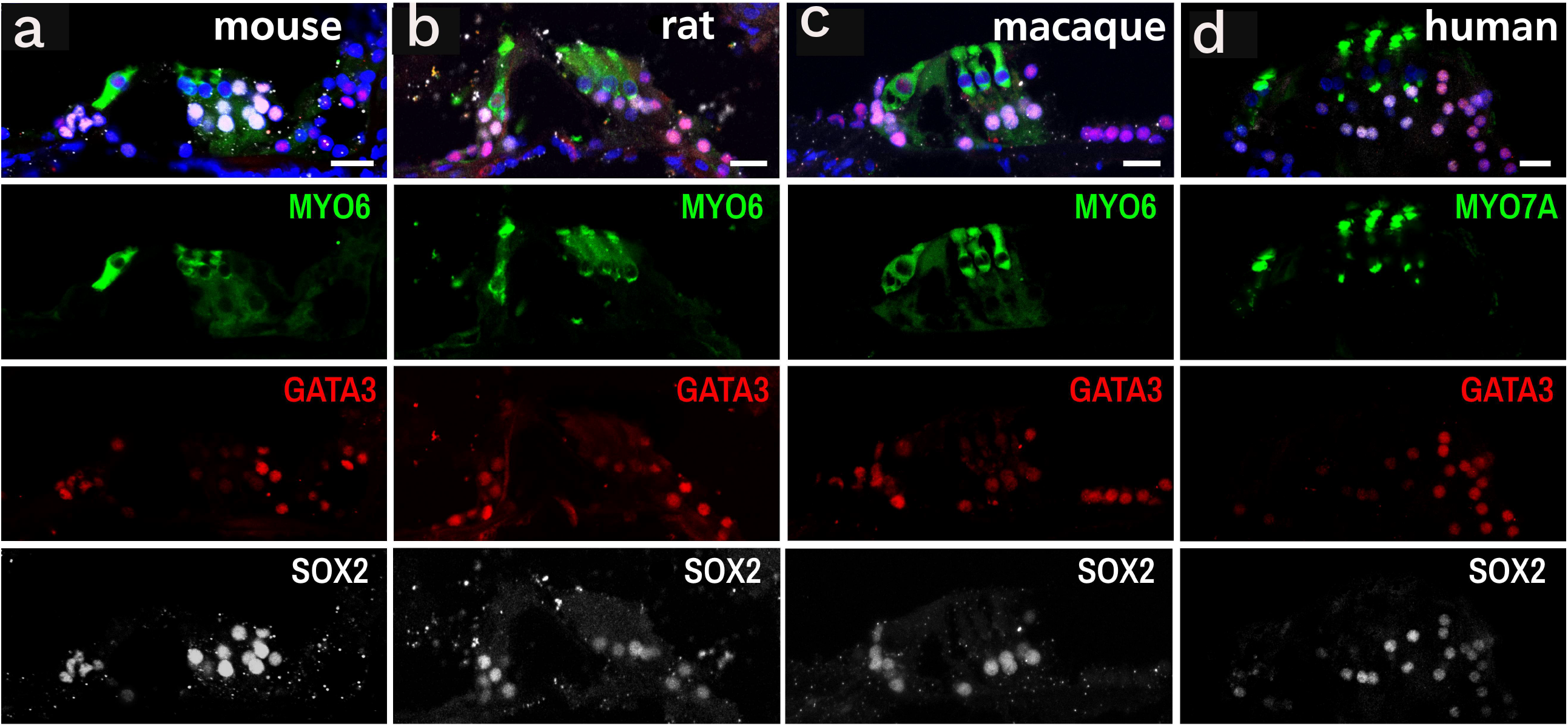
Cross-species comparison of potential *GATA3* regulatory elements identified from FANTOM5 human CAGE-seq data. (a) Putative promoter sequences identified ±100kb from the TSS of *GATA3* on human chromosome 10 were compared for sequence similarity against corresponding sequences in the macaque, rat, and mouse genomes. (b) Putative GATA3 enhancer sequences identified on human chromosome 10 were compared to corresponding sequences in the macaque, rat, and mouse genomes. Heat maps use purple-yellow-green to reflect the percent similarity of the indicated sequences ranging from 40 – 100% identity (shown as 0.4-1.00). Red boxes indicate sequences that have been previously identified in the literature and validated as regulators of *GATA3* or *Gata3* expression (see supplemental table S1). Blue boxes indicate sequences identified as likely important in sensory epithelial cells (SEC) or neural stem cells (NSC). The black box indicates the only sequence identified in this analysis that appears to be unconserved in macaques, rat, and mice as compared to the human sequence. *A sequence identified in Gregoire and Romeo 1999 as being non-conserved between humans and mice. (c) Comparison of amino-acid sequences for GATA3 across humans, macaques, rats, and mice suggests a very high level of conservation at the peptide level. (d) Image representing the amino acid sequence of GATA3 protein in human, macaque, rat and mouse recognized by the R&D and BD-Bioscience anti-GATA3 antibodies used for this study.

### Image processing

All the images were imported to and processed by Image J software. Far red channel 647 was split from the other channels and the following despeckling processing was performed for the 647 channel only. Open image> adjust brightness and contrast>Process>Noise> Remove outlier > process> sharpen. Image J software was also used to obtain mean gray values as a measure of fluorescence intensity. In unadjusted images, an ROI was generated to encircle GATA3 immunolabeled nuclei and the mean gray values were measured (ctrl+M). After each measurement, the ROI was moved adjacent to the positive nucleus to obtain a measure of background fluorescence. At least 4 nuclei from PCs and DCs were recorded and at least 4 from Hensen’s, Claudius, and Inner Phalangeal cells were recorded from each image. At least one image from each individual was measured, and a sample of N= 4, 2 male and 2 female, was used for each species in this comparison (4 mice and 4 humans). Background fluorescence measures were subtracted from nuclear measures and these normalized values for PCs and DCs were compared to the normalized values from Hensen’s and Claudius cells using two-tailed, dependent measures t-tests.

## RESULTS

### Cross species comparison of potential GATA3 regulatory sequences

We used SlideBase to query the Fantom5 database which contains CAGE-seq (cap analysis gene expression sequencing) data from primary human tissue from several different organs as well as hundreds of cell lines. From this, we identified potential promoters and enhancers of *GATA3* that were located within 100 kilobases in either direction of the *GATA3* transcriptional start site. This search yielded 32 potential promoters and 10 potential enhancer or repressor elements (Figure 1, Supplemental data table 1). The orthologues of *GATA3* in the crab eating macaque, Norway rat, and house mouse reside in chromosomes 9, 17, and 2, respectively. The sequences from these chromosomes were queried to identify the corresponding promoter or enhancer sequences and the degree of homology was computed. Figure 1 illustrates that all of the identified regulatory elements, with the exception of one potential promoter sequence, are highly conserved between the human and macaque genomes. In contrast, a significant portion of the identified regulatory elements were less than 75% conserved between the rodent species and humans. For the potential promoters, sequences located closer to the GATA3 coding region were more likely to show greater homology between rodents and humans while sequences that were more distal in the 5’ direction were less conserved.

We also conducted a literature search to find GATA3 regulatory elements that have been previously identified and validated in either mouse or human cells (Supplemental data table 1) and several of these matched or overlapped with sequences obtained from the Fantom5 data suggesting the utility and validity of the Fantom5 CAGE-seq data for identifying putative regulatory sequences. Fantom5 data also provides annotation of the potential promoter and enhancer sequences based on the cell or tissue types from which the greatest numbers of mapping-reads were obtained. One of the potential promoter sequences was identified as highly active in sensory epithelial cells, and two of the enhancers were identified as active in neuronal stem cells (Figure 1, Supplemental Table 1), suggesting that these three regulatory elements may be important in regulating GATA3 expression in inner ear sensory cells and developing SGNs. Two of these three sequences were fairly highly conserved across all four species, however one of the enhancer regions identified as being prevalent in neural stem cells was not very well conserved between rodents and primates. Given the number of different cells and tissue types in which GATA3 is expressed, it is not surprising that numerous potential promoters and enhancers were identified and that some were conserved while others were not. However, the presence of both conserved and non-conserved regulatory elements leaves open the question of whether the distribution of GATA3 might be conserved or not, specifically within the inner ear tissues of these species. Thus, we mapped the immunocytochemical distribution of GATA3 in the adult ears of mice, rats, macaques and humans.

### Validation of GATA3 immunostaining

The peptide sequence for GATA3 is very highly conserved across mouse, rat, macaque, and human species (Supplemental figure S1) (figure 1d). Commercial antibodies against GATA3 were obtained and expected to have binding affinity for the protein in all 4 species, and one in particular (BD Biosciences cat#558686) was selected due to being raised against a region of the peptide sequence that is 100% conserved across all 4 species (Supplemental figure S1) (figure 1d). To validate GATA3 immunostaining, several approaches were taken.

First, neonatal mouse cochleae were immunostained to confirm the previously published pattern of expression in mice at this age (Nishimura et al., 2017; Walters et al., 2017). Indeed, robust nuclear staining was readily detectable in all cochlear SCs and in the vast majority of SGNs (Figure 2) which is consistent with previous reports. Next, we examined human tissues which could be used as positive controls including human tonsil (Atayar et al., 2005) and human spiral ganglion tissue from a young individual (8 year old male). We also ran absorption controls on these tissues by pre-incubating the GATA3 antibodies with a recombinant human GATA3 peptide, as well as a secondary-only control. GATA3 immunoreactivity was readily detectable in human tonsil samples as well as in the SGNs from an 8 year old male and immunoreactivity in human tissues was abolished when the antibodies were pre-absorbed by GATA3 peptide (Figure 2). Controls where only the secondary antibody was used were also negative for any detectable nuclear staining. We also immunostained inner ear tissues from a *Gata3* conditional knockout mouse (Grote et al., 2008) that was bred with an Fbxo2CreER mouse line (Hartman et al., 2018) to target the cells of the cochlear sensory epithelium. Following tamoxifen induction, GATA3 immunoreactivity was abolished in the Cre-positive cells of these mice (Figure 2). Thus the GATA3 antibodies used in this study demonstrated specificity for GATA3 in both mouse and human tissues.

**Figure 2:**
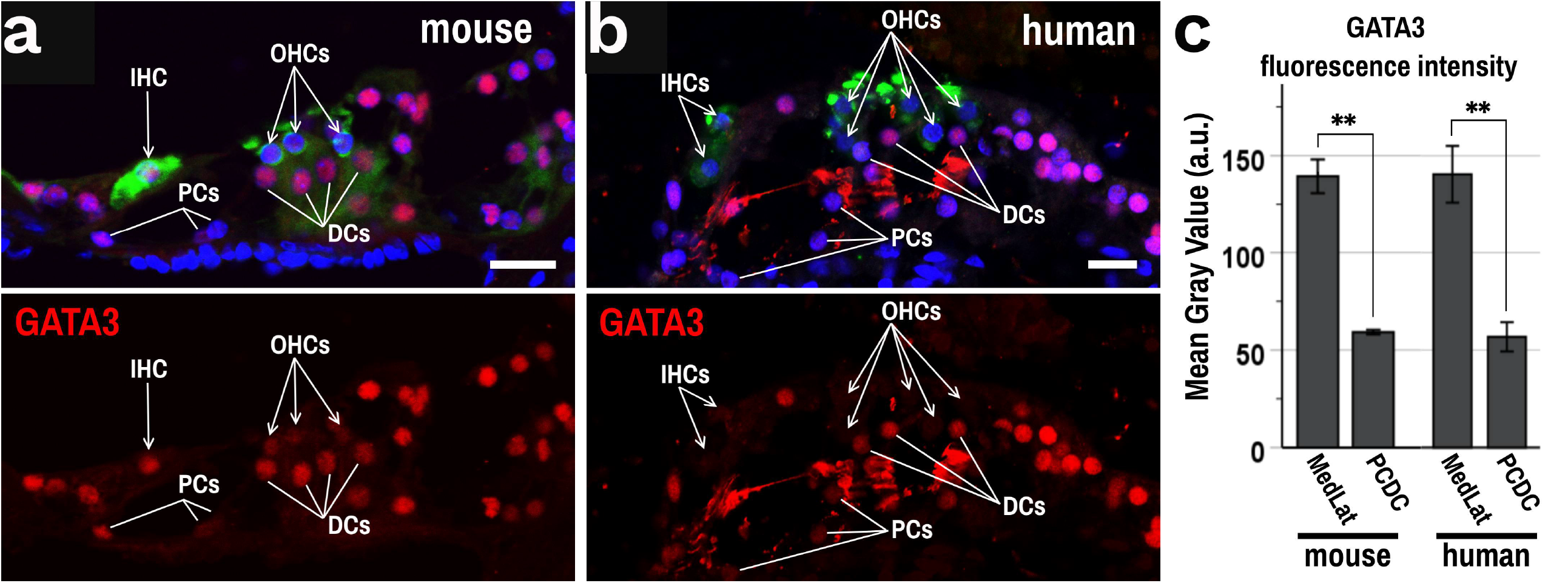
Validation of GATA3 antibodies in mice and humans. (a) A graphical representation of organ of Corti, showing the anatomical location of outer hair cells (OHC), inner hair cells (IHC) and supporting cells (SCs) such as Dieter’s cells and Pillar cells (b, b’). Immunostaining with GATA3 antibody (BD biosciences) in mice at postnatal day 0 (P0) replicated what has been previously reported. (b) GATA3 (magenta) is readily detectable in the nuclei of all cochlear SCs as well as in IHCs, though only faintly detectable at this point in OHCs. Hair cells labeled using myosinVIIa (MYO7A) antibody (green). (c) Cryosection of neonatal mice temporal bone showing apical to basal gradient of GATA3 (magenta) expression in NF-H (green) SGN cell bodies. (c’ and c’’) Zoomed in images (from c) of GATA3 expression from the apex shows at this age, most of SGN cell bodies express GATA3. (d) GATA3 immunostaining (red) is robust in cells of the cochlear outer sulcus (OS) in adult mice. In Fbxo2CreER+;Gata3(loxp/loxp) mice that were not induced with tamoxifen, this robust GATA3 immunoreactivity persists and can be clearly seen in OS cells that also express the Venus reporter driven by the Fbxo2 promoter. (e) Conditional deletion of *Gata3* using the Fbxo2CreER and tamoxifen induction at 6 weeks of age abolishes GATA3 immunoreactivity in many Venus positive OS cells. Hoechst dye (HXT, blue) was used to counterstain nuclei. (f) GATA3 immunostaining using diaminobenzidine (DAB) color reaction shows clear brown color reaction in the nuclei of lymphocytes in human tonsil samples, a known positive control tissue for GATA3 immunostaining. (g) GATA3 immunoreactivity with DAB was also detectable in the nuclei of human SGNs. (h) However, if the GATA3 antibody was pre-absorbed with GATA3 peptide, then nuclear GATA immunolabeling could be abolished. (i) Similarly, applying secondary antibodies only to human SGN sections did not reveal any nuclear DAB staining. Scale bars = 20 μm.

### GATA3 distribution in the mature Organ of Corti

In adult mice, rats, macaques, and humans, GATA3 was readily detectable in the non-sensory SCs medial and lateral to the organ of Corti (Figure 3). Specifically, inner border and inner phalangeal cells (IPhcs) medial to the IHCs, and Hensen’s and Claudius cells lateral to the OHCs were strongly immunopositive for GATA3. In these four species, PCs and DCs did not exhibit robust GATA3 immunoreactivity. When gain was significantly increased on the confocal microscope so as to cause oversaturation in the SCs medial and lateral to the HCs, nuclear GATA3 immunostaining could be detected in PC and DC nuclei suggesting that GATA3 is still expressed in adult PCs and DCs, albeit at levels much reduced compared to IPhcs, Hensen’s cells, and Claudius cells (Figure 4). Quantification and comparison of mean gray values of fluorescence intensity confirmed that GATA3 immunostaining was significantly brighter in cells medial and lateral to the HCs, such as Hensen’s and Claudius cells, compared to PCs and DCs (Figure 4). Indeed this pattern held true for both mouse (t(3) = 10.5, p = 0.002) and human (t(3) = 7.0, p = 0.006) cochleae.

**Figure 3:**
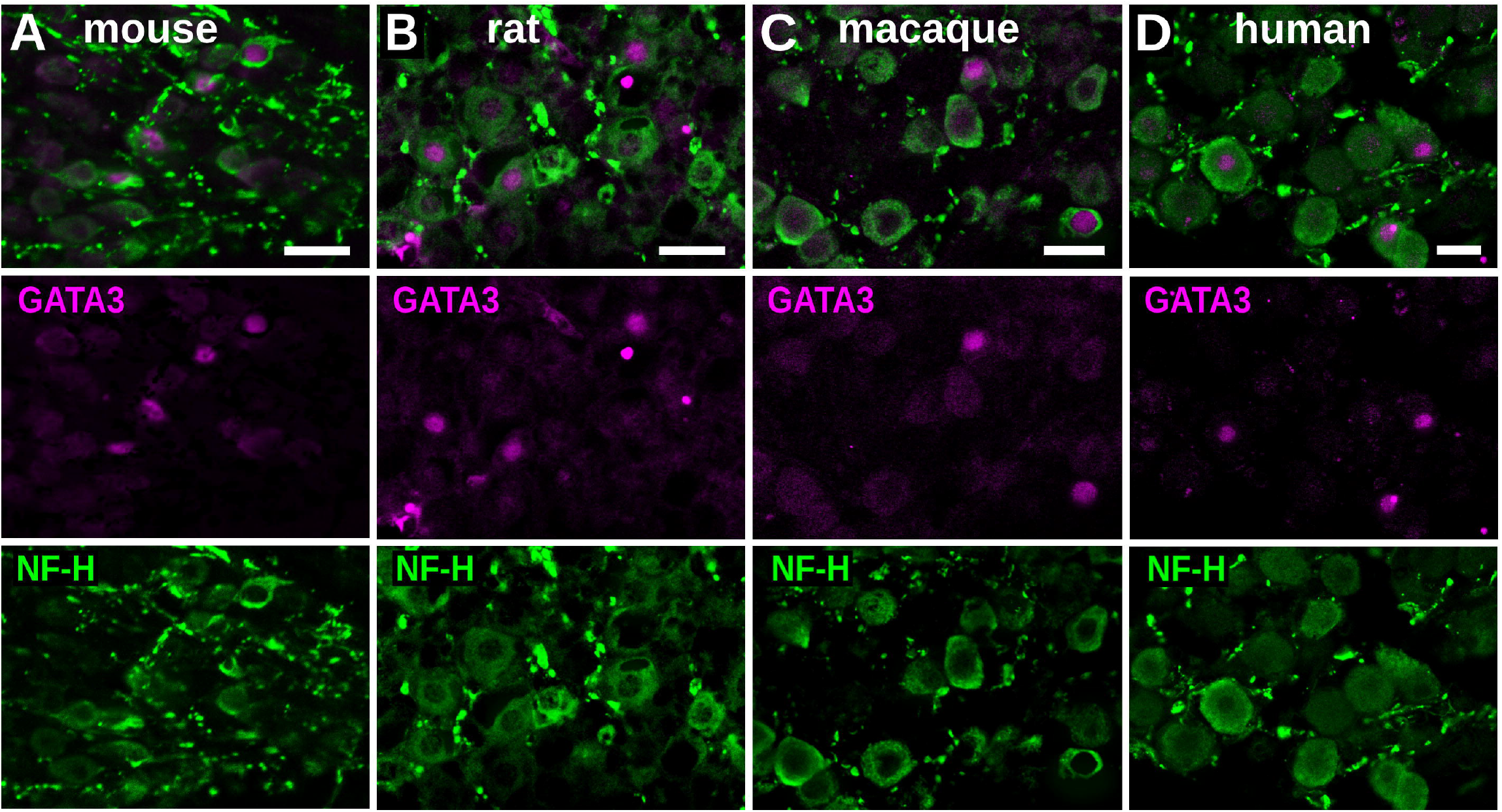
GATA3 expression in the organ of Corti across species. Cryosections of the organ of Corti from different species including adult mouse (a), rat (b), macaque (c) and human (d) were immunolabelled for HC markers Myo6 or Myo7A (green), SC marker Sox2 (white), and GATA3 (red). The nuclei were stained with Hoechst (blue). Strong GATA3 labelling was found to be in the SCs medial and lateral to the PCs and DCs including Inner phalangeal, Claudius, and Hensen’s cells in all the species. IHCs were also GATA3 positive in all species *except humans* where GATA3 was not detected in IHCs in any samples. GATA3 labeling in the nuclei of PCs and DCs of all species was much fainter than in other SCs and inconsistently detected. In all 4 of the species examined, GATA3 was largely undetectable in the OHCs. For each species, N ≥ 4, Scale bar = 20μm.

**Figure 4:**
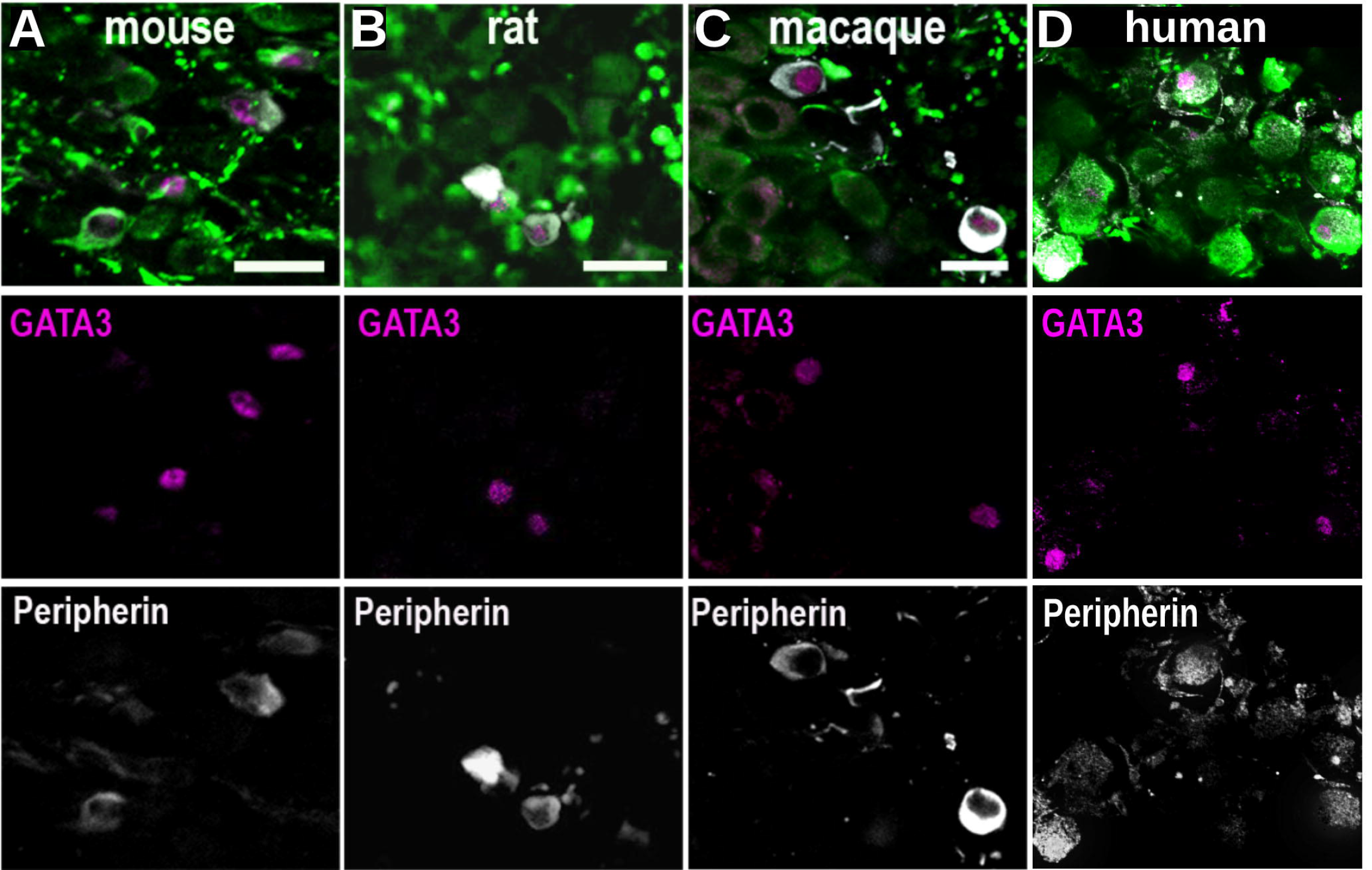
GATA3 immunofluorescence has significantly greater optical intensity in Hensen and Claudius cells than in Pillar and Deiters’ cells. (a-b) Oversaturated images of mouse (a) and human (b) organs of Corti reveal the presence of GATA3 in PCs and DCs. (c) In images that were not oversaturated, quantification of mean gray values (normalized to background) confirmed that GATA3 immunostaining was significantly brighter in cells medial and lateral (MedLat) to the HCs, such as Inner phalangeal cells and Hensen’s and Claudius cells, compared to PCs and DCs This pattern was true for both mouse (t(3) = 10.5, p = 0.002) and human (t(3) = 7.0, p = 0.006) cochleae. Scale bar = 20μm.

In mouse, rat and macaque, GATA3 was detectable in the IHCs and was largely undetectable in the OHCs (Figure 3). However, in some sections, faint GATA3 nuclear staining could be occasionally seen in OHCs when confocal gain and image brightness were significantly increased (Figure 4), though this was not consistently observed. This phenomenon was also not observed in any human cochleae. Therefore, while the extent to which GATA3 may be expressed in adult cochlear OHCs may require further investigation, it is clearly significantly diminished compared to the surrounding cells. This pattern is consistent with what has been previously published in a mixed background strain of mice and is replicated here in an inbred CD1 strain. However, it is of note that, while IHCs in mice, rats, and macaques, all demonstrated GATA3 immunoreactivity, GATA3 was not detected in the inner HCs in any of the human temporal bone samples regardless of which GATA3 antibody was used. There was no detectable GATA3 immunostaining in the vestibular ganglia nor in the hair cells or SCs of the vestibular organs in any of the species examined (data not shown).

### GATA3 distribution in the Spiral Ganglion

It has been previously reported in mice that GATA3 immunoreactivity, while readily detectable in all of the developing SGNs, diminishes to undetectability in the type I SGNs as the mice age. Here, we used GATA3 antibodies from two different vendors (BD Bioscience and Santa Cruz) to determine GATA3 expression in the SGNs in not only mouse, but also rat, macaque, and human cochleae. In newborn (P0) mice, strong GATA3 labelling was observed in all NF-H positive neuronal nuclei (Figure 2). On the contrary, in adult mice, as well as adult rats, macaques, and humans, GATA3 immunoreactivity was restricted to only a subset of NF-H positive SGNs (Figure 5). As has been previously reported in mice, where GATA3 is believed to be highly enriched in peripherin-expressing type II SGNs (Nishimura et al., 2017), GATA3-positive neurons in rat, macaque, and human samples also appeared to be generally co-labeled by peripherin antibodies (Figure 6). This co-labeling with peripherin suggests that GATA3 is predominantly expressed in type II SGNs in adults of all four species.

**Figure 5:**
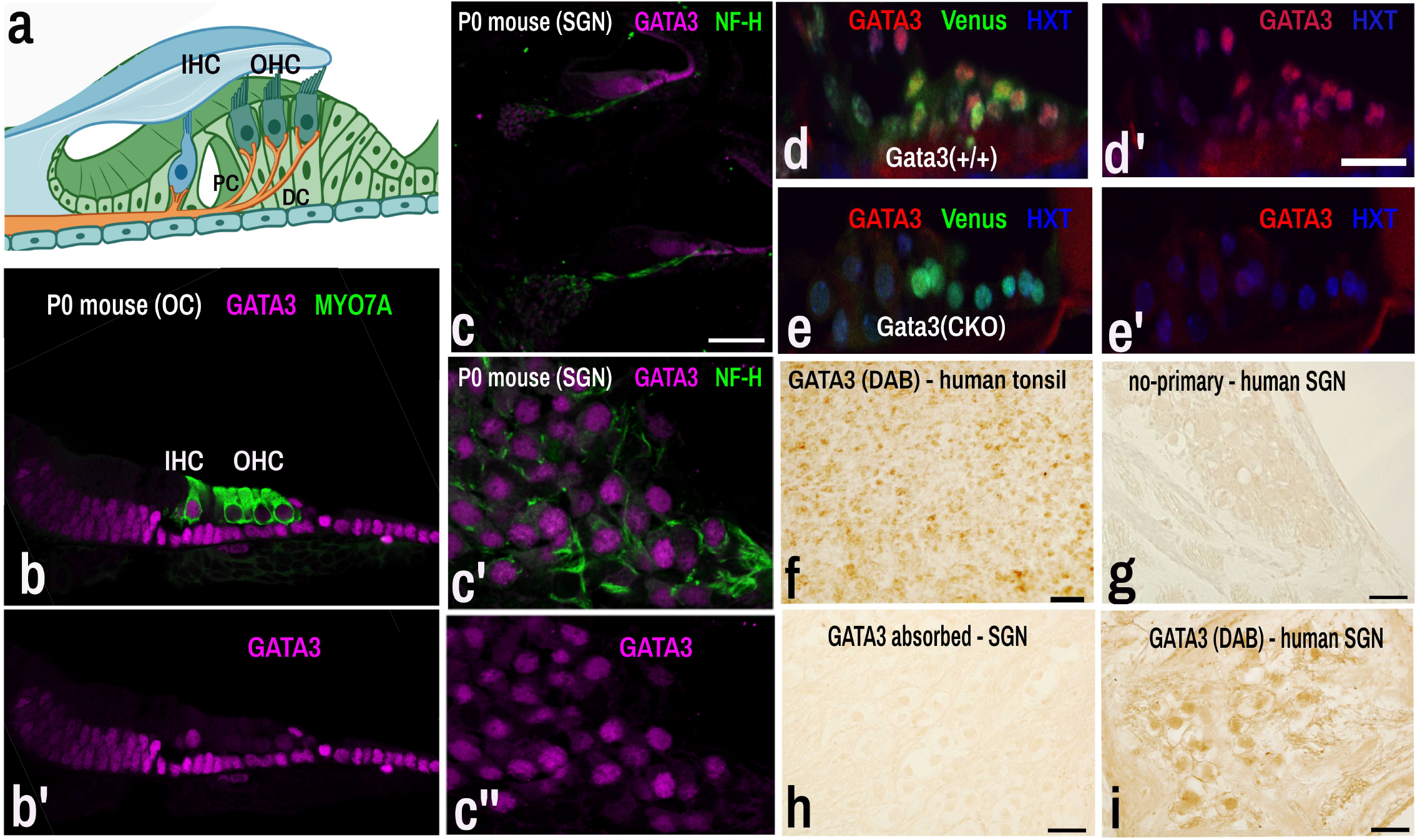
GATA3 expression in the spiral ganglion. Cryosections of SGN from mouse (a), rat (b), macaque (c) and human (d) samples were immunolabeled with neuronal marker NF-H (green) and GATA3 (magenta) antibodies (BD Bioscience). In adult mice, as well as adult rats, macaques, and humans, GATA3 immunoreactivity was restricted to only a subset of NF-H positive SGNs (Scale bar = 20μm).

**Figure 6:**
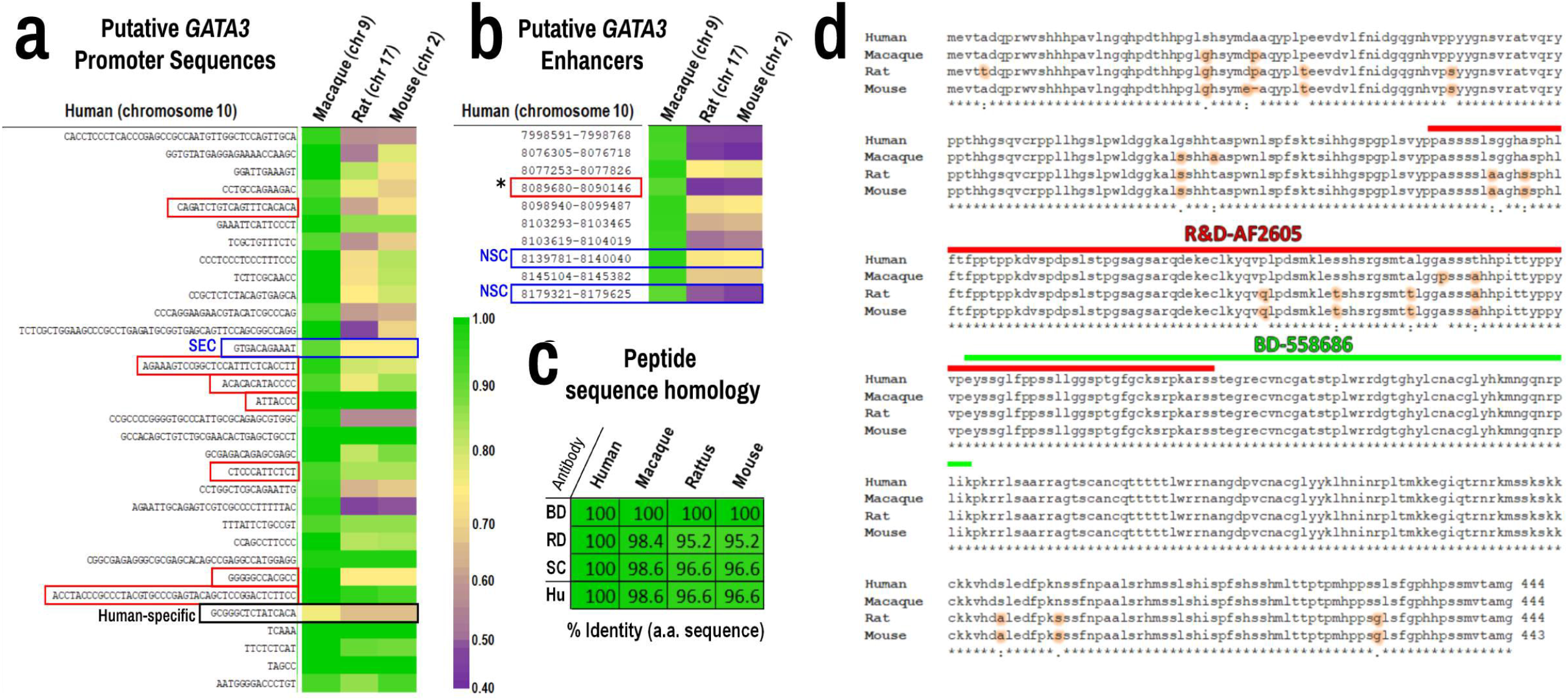
GATA3 and peripherin co-labeling suggest GATA3 is expressed in type II SGNs. Peripherin immunolabeling has traditionally been used to distinguish type II SGNs from type I SGNs. In mice (a), rats (b), macaques (c), and humans (d) GATA3 (magenta) largely co-labeled peripherin (white) positive cells suggesting GATA3 is mostly present in type II SGNs. NF-H (green) was used to label all SGNs, type I and type II. Scale bar = 20 μm.

## DISCUSSION

GATA3 is critically important in the development and maintenance of a number of tissues and cell types. This is particularly true in the case of the auditory periphery where both *GATA3* haploinsufficiency in humans and *Gata3* deletion from mice reveal that it is essential for cochlear development and function. Recently, it was shown in mice that the downregulation of GATA3 from cochlear SCs with age may be a key factor that prevents the regeneration of cochlear sensory hair cells. Indeed, several recent studies have suggested that GATA3 may be a universal mediator of injury-induced regeneration across several tissue types (Kizil et al., 2012; Liao et al., 2017; Montserrat et al., 2013; Strühle & Schmidt, 2012). In the mouse inner ear, specifically, it has been shown that reinstating *Gata3* expression in adult PCs and DCs can enable these cells to better respond to ectopic *Atoh1* and be converted into HC-like cells (Walters et al., 2017). The importance of such regeneration studies lies in the hopes of translating the findings from mice or other laboratory animals to humans, where a therapy to regenerate cochlear HCs could potentially lead to a rehabilitation of hearing ability for the vast number of individuals who suffer from sensorineural hearing loss. However, recent reviews from multiple fields have demonstrated that a worrying number of studies in rodents fail to translate to effective therapies in humans (Hackam & Redelmeier, 2006; Kuroda et al., 1999; Lees et al., 2006; Marshall et al., 2001; Sydserff et al., 2002). Among other reasons contributing this, one fact could be the expression of even the highly conserved genes between rodents and human, are regulated by different regulatory elements. It results in differential spatiotemporal regulation and functions of orthologous genes. This may be due to a number of causes, not least of which is the fact that, even when the open reading frames of genes are highly conserved between rodents and humans, the regulatory elements, the temporospatial patterns of expression, and even the functions of these orthologs may not be so well conserved. Indeed, several studies from the field of inner ear biology have suggested that genes critical for hearing in humans do not always have a conserved role in the hearing of mice (Liu, 2000; Plum et al., 2001; Xia et al., 1998). Other studies have shown that proteins that are critical for hearing function in mice, do not exhibit the same patterns of expression in the inner ears of primates (Hosoya, Fujioka, Ogawa, et al., 2016; Hosoya, Fujioka, Okano, et al., 2016; Kim et al., 2014; Matsuzaki et al., 2018; Suzuki et al., 2016). Thus there is a critical need to better understand the potential similarities or differences in the regulation and expression of genes and proteins known to be vital to hearing function, especially when those genes are targets for potential human therapies such as ATOH1.

As stated above, GATA3 appears to be critical for HC regeneration approaches that rely on ATOH1. This would include not just ectopic expression of *Atoh1*, but also Notch inhibition which is presumed to operate mechanistically via *Atoh1* upregulation (Mizutari et al., 2013). Critically, clinical trials aimed at regenerating HCs have begun for both *ATOH1* gene therapy and pharmacological inhibition of Notch signaling. Thus, knowing whether GATA3 is reduced or absent from cochlear SCs in human ears, as it is in mice, can provide some insight into how effective these approaches may be. Here we examined the expression of GATA3 in adult mice, rats, macaques and humans and found that expression patterns in SCs and SGNs was conserved. Specifically, PCs and DCs do not exhibit robust GATA3 immunolabel like their neighboring SCs medial and lateral to the organ of Corti. In the SGNs, type II SGNs remain GATA3 positive into adulthood, and likely throughout life. With regard to SCs and *ATOH1*-mediated therapeutic approaches, this suggests that what was found in the mouse model (i.e. that *Gata3* may need to be reinstated to obtain a robust effect from *Atoh1*), likely also holds true for humans. As such, if current clinical trials that target *ATOH1* or Notch should deliver lackluster results, then perhaps combined targeting of *GATA3* and *ATOH1* may prove beneficial.

With regard to SGNs, the functions of GATA3 are well studied during inner ear development. However, the function of GATA3 in the adult cochlea are not as well-known, but there is some evidence to suggest that, in mice, the persistent expression of high levels of GATA3 is necessary for SGN survival (Hoshino et al., 2019). Haploinsufficiency of *Gata3* in mice leads to apoptotic loss of SGNs first seen at around 5 months of age. Our data suggest that in human type II SGNs retain the expression of GATA3 in the adulthood, and this conservation and preservation of GATA3 expression GATA3 may contribute to the survival and function of these SGNs. Indeed, while it has been shown that SGNs are lost with age in humans (Schuknecht & Gacek, 1993; Schuknecht, 1964), some evidence suggests that type II SGNs may survive the aging process better than type I SGNs (Spoendlin & Schrott, 1988). However, such a survival advantage for type II SGNs has been contested (Zimmermann et al., 1995).

Though more recent evidence suggests that, while type I SGNs may not die off in greater numbers, perhaps the type I auditory nerve fibers, and their connections to IHCs, could be more susceptible to degeneration with age (Wu et al., 2019). While the extent to which such survival or innervation differences occur with age in humans remains unresolved, the similar expression patterns of GATA3 in the adult SGNs of mice, rats, macaques, and humans, suggests that rodents and non-human primates may provide adequate models for better understanding the role of GATA3 in SGN survival and function in the adult human cochlea.

Another interesting finding here was that the pattern of GATA3 expression, while largely similar, was not identical across all four species. Indeed, in mice, rats, and macaques, cochlear IHCs were GATA3 positive, but GATA3 was not observed in the IHCs of any of the human samples studied (with ears from at least 12 different individuals having been examined).

Furthermore, while faint GATA3 immunostaining was occasionally seen in mouse, rat, and macaque OHCs when confocal gain and image brightness were significantly increased, such increases did not reveal any hints of GATA3 expression in either IHCs or OHCs in human cochlear sections (Figure 4b). If GATA3 expression is differentially regulated in human cochlear HCs compared to these other species, this could have potential implications for understanding the functions of GATA3 in the adult human cochlea.

GATA3 is known to play important roles in sensory cell survival and in the formation and maintenance of synapses (Hoshino et al., 2019; Yu et al., 2013). As noted above, neural degeneration is a demonstrated problem of aging within the human ear and is suspected to contribute to presbycusis (Nadol, 1990; H. F. Schuknecht & Gacek, 1993; Wu et al., 2019). Thus, the relative absence of GATA3 from IHCs in humans could be a contributing factor to the losses of auditory nerve fibers and SGNs and suggest GATA3 as a potential therapeutic target for mitigating age-related degeneration of the auditory nerve. However, given that this may be a phenomenon specific to human cochleae, experimental models other than traditional rodent or non-human primate models may have to be developed to directly test this hypothesis. One such approach to pursue this could be through the identification of the genetic regulatory elements that lead to the absence of GATA3 in human cochlear hair cells. Here we identified dozens of potential regulatory elements, including several that are not conserved between humans and rodents. Close examination of the patterns of homology for these elements across the species examined can therefore help to generate hypotheses regarding which elements may be involved. For example, our analysis of human CAGE-seq data (Supplemental Table S1) confirms findings from a previous study in identifying a GATA3 regulatory element that is present in human sequence, but not in mouse or rat (Grégoire & Roméo, 1999). However, this particular regulatory element is largely conserved in the macaque genome and so is less likely to be responsible for the human-specific pattern of GATA3 absence from IHCs. Of the putative regulatory sequences we examined, one sequence was identified that did not show homology between macaques and humans, but was conserved between rodents and macaques (Figure 1). This potential regulatory element, therefore, is an ideal candidate for us to more closely investigate in future experiments to determine whether it may be involved in the human-specific pattern of GATA3 expression, or lack thereof, in the IHCs.

One potential avenue for testing putative human-specific regulatory elements could be via the introduction of these sequences in mice by CRISPR-mediated gene editing or massive parallel reporter assay to determine whether GATA3 expression is subsequently abolished from IHCs in adult mice cochleae (Lambert et al., 2021; K. Li et al., 2020). Following this, the effects of such a loss of GATA3 on IHC or SGN function and survival can be assayed. However, it is important to note that our study of human tissues was limited by the fact that the donors were largely of advanced age at the time of death, whereas the rodents and macaques were of more middling ages (5 months old and 4-6 years old, respectively). While matching corresponding ages to an exact degree is difficult across species with differing lifespans, it is certainly plausible to consider the human subjects as being relatively older than the corresponding ages of the rodents and macaques. However, regardless of whether the inter-species difference in IHC expression of GATA3 is an artifact of aging differences or represents an accurate indication of differential GATA3 regulation and distribution, the findings here are novel and faithfully describe a lack of GATA3 in adult human IHCs particularly in comparison to the robust expression seen in neighboring cells. This finding reinforces the importance of additional study into the function of GATA3 in adult IHCs in both animal and human models.

Taken together, the data here suggest that, there are many putative regulatory regions for GATA3, some well-conserved between primates and rodents, and others that are not as well-conserved. Also the findings suggest that the pattern of GATA3 expression is largely similar in the inner ear across mice, rats, macaques, and humans, particularly for cochlear SCs and SGNs. Importantly, however, human IHCs appear to lack GATA3, and regardless of whether this is due to advanced age, non-conserved regulatory elements, or both, it suggests an intriguing difference between rodent models at the ages they are traditionally studied and aged humans. Overall, however, these findings provide greater confidence that data pertaining to GATA3 in SGNs and SCs in rodents may, for the most part, similarly hold true in humans, and therefore may inform regenerative therapeutic strategies. However, they also suggest potential regulatory regions worthy of further investigation, particularly with regard to the regulation and expression of GATA3 in cochlear HCs.

## Acknowledgements

The authors would like to thank Drs. Marianne Conway and Al Sinning for their assistance as directors of the UMMC body donation program, and of course, a tremendous thanks to the individuals who generously donated their corporeal selves or that of their loved ones to the UMMC body donation program and to the NIDCD National Temporal Bone Laboratory at UCLA. Their generosity has helped to advance both medical education and scientific research progress. Also, our thanks go to the Center for Comparative Research at UMMC for assistance with animal care and husbandry, and Drs. Parminder Vig and G. Lee Bidwell for the use of their confocal microscope. Thanks to Drs. Paul May and Susan Warren who provided macaque temporal bones for use in this study, and thanks to Dr. Bernadette Grayson who provided rat temporal bones for this study.

