## Supplementary figures and images for "Identification of putative GATA3 regulatory elements and comparison of GATA3 distribution in cochleae of mice, rats, macaques, and humans"

### Supp_Figure_S1_GATA3 -min.jpg

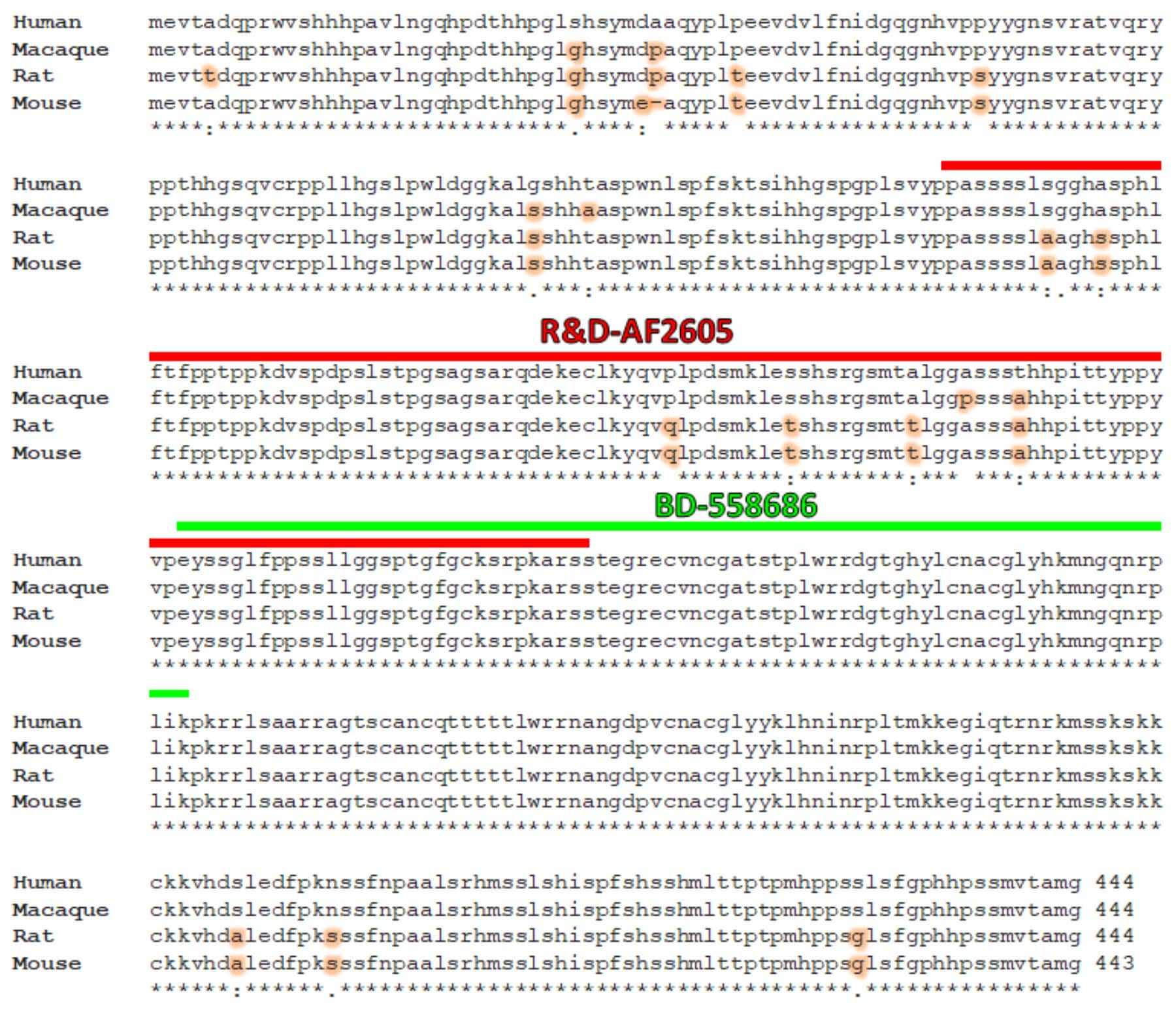
